# Toward defining deep brain stimulation targets in MNI space: A subcortical atlas based on multimodal MRI, histology and structural connectivity

**DOI:** 10.1101/062851

**Authors:** Siobhan Ewert, Philip Plettig, M. Mallar Chakravarty, Andrea Kühn, Andreas Horn

## Abstract

Three-dimensional atlases of subcortical brain structures are valuable tools to reference anatomy in neuroscience and neurology. In the special case of deep brain stimulation (DBS), the three most common targets are the subthalamic nucleus (STN), the internal part of the pallidum (GPi) and the ventral intermediate nucleus of the thalamus (VIM). With the help of atlases that define the position and shape of these target regions within a well-defined stereotactic space, their spatial relationship to implanted deep brain stimulation (DBS) electrodes may be determined.

Here we present a composite atlas based on manual segmentations of a multi-modal high-resolution MNI template series, histology and structural connectivity. To attain exact congruence to the template anatomy, key structures were defined using all four modalities of the template simultaneously. In a first step tissue probability maps were defined based on the multimodal intensity profile of each structure. These observer-independent probability maps provided an excellent basis for the subsequent manual segmentation particularly when defining the outline of the target regions.

Second, the key structures were used as an anchor point to coregister a histology based atlas into standard space. Finally, a sub-segmentation of the subthalamic nucleus into three functional zones was estimated based on structural connectivity. The resulting composite atlas uses the spatial information of the MNI template for DBS key structures that are visible on the template itself. For remaining structures, it relies on histology or structural connectivity. In this way the final atlas combines the anatomical detail of a histology based atlas with the spatial accuracy of key structures in relationship to the template anatomy. Thus, the atlas provides an ideal tool for the analysis of DBS electrode placement.

**Highlights:**

- Composite subcortical atlas based on a multimodal, high definition MNI template series, histology and tractography
- High definition atlas of DBS targets exactly matching MNI 152 NLIN 2009b space
- Multimodal subcortical segmentation algorithm applied to MNI template

## Introduction

Three-dimensional subcortical atlases are valuable tools to reference anatomy in the brain. In the field of deep brain stimulation (DBS), atlases of certain target structures may be used to study the relationship of electrode placement to its target structure in subcortical space (Merkl et al. 2015; Horn & Kühn 2015; Neumann, Jha, et al. 2015; Neumann, Staub, et al. 2015; Eisenstein et al. 2014; Barow et al. 2014; Welter et al. 2014; Butson et al. 2007). Most of the available atlases have been defined either by histology or magnetic resonance imaging (MRI). For instance, the atlases by Yelnik et al. (Yelnik et al. 2007) and Chakravarty et al. (Chakravarty et al. 2006) were defined using histological stacks of a single brain. Mai et al. defined a histological atlas based on three brains (Mai et al. 2007) and Morel et al. used maps derived from six brains (Morel 2013). In case of the Chakravarty study, the atlas was coregistered to an MRI template by intensity-matching of segmented histological sub-structures to the same structures defined by the template. This led to intensity-matched *pseudo-MRIs* that could be nonlinearly registered to a standard template. In this fashion, the atlas was transformed into common space (MNI *colin27*) to allow for conclusions in standardized brain anatomy. Other subcortical atlases were derived based on MRI. Briefly, the BGHAT atlas was defined by manual delineation of subcortical structures on a single MRI image (Prodoehl et al. 2008). Similarly, the CFA subcortical shape atlas was manually delineated on MRI, but used acquisitions of 41 subjects (Qiu et al. 2010). The ATAG atlas used multimodal high field MRI to estimate structures of the basal ganglia on a series of images (Keuken et al. 2014) and even included three probabilistic maps of the subthalamic nucleus (STN) based on cohorts with varying age (Keuken et al. 2013). In a study by Lenglet et al., a comprehensive atlas of basal ganglia structures and their white-matter interconnections was established using structural and diffusion-weighted MRI (Lenglet et al. 2012). Finally, the MIDA model (Iacono et al. 2015) focused on a detailed whole-head segmentation that included structures like blood vessels, muscles and bones but also covered subcortical structures of the brain including the STN.

However, despite the high precision of anatomical labels in histological atlases and the probabilistic nature of atlases derived from larger cohorts of subjects, a shortcoming of the aforementioned atlases is that their spatial definition of structures – such as the STN – do not exactly match the position, size and shape of the same structures defined by the MNI template (Fig. 1). This is a crucial point since it means that studies using these atlases would use the MNI templates (for nonlinear warps) in combination with descriptions of structures not exactly matching the template. Moreover, this problem is further aggravated in the field of DBS imaging where i) anatomical structures are small in size and ii) a shift of a few millimeters can significantly alter clinical outcome. Any standardized stereotactic space is defined solely by its anatomical template(s). In other words, these templates define which anatomical structure is located at any given Cartesian coordinate. The STN is a good example to further illustrate the issue. The nucleus is well visible as a hypointense lentiform structure on T2-weighted versions of, for instance, the ICBM 152 2009b nonlinear asymmetric template (Fig. 1, upper left panel). If used in a nonlinear normalization process, this means that any algorithm will try to warp the visible parts of the nucleus from single subject anatomy onto the lentiform structure defined by the template. In theory, a perfect algorithm would even obtain a solution that renders a patient's STN equally lentiform, equally sized and placed at the exact same spatial position as the one in the template. The better the algorithm, the better it will achieve this goal (Klein et al. 2009; but see discussion). Contrastingly, STN parcellations of available atlases do not necessarily match the size, position and shape of the hypointense area that marks the STN inside the MNI templates (Fig. 1; see S1 for details).

**Figure 1:**
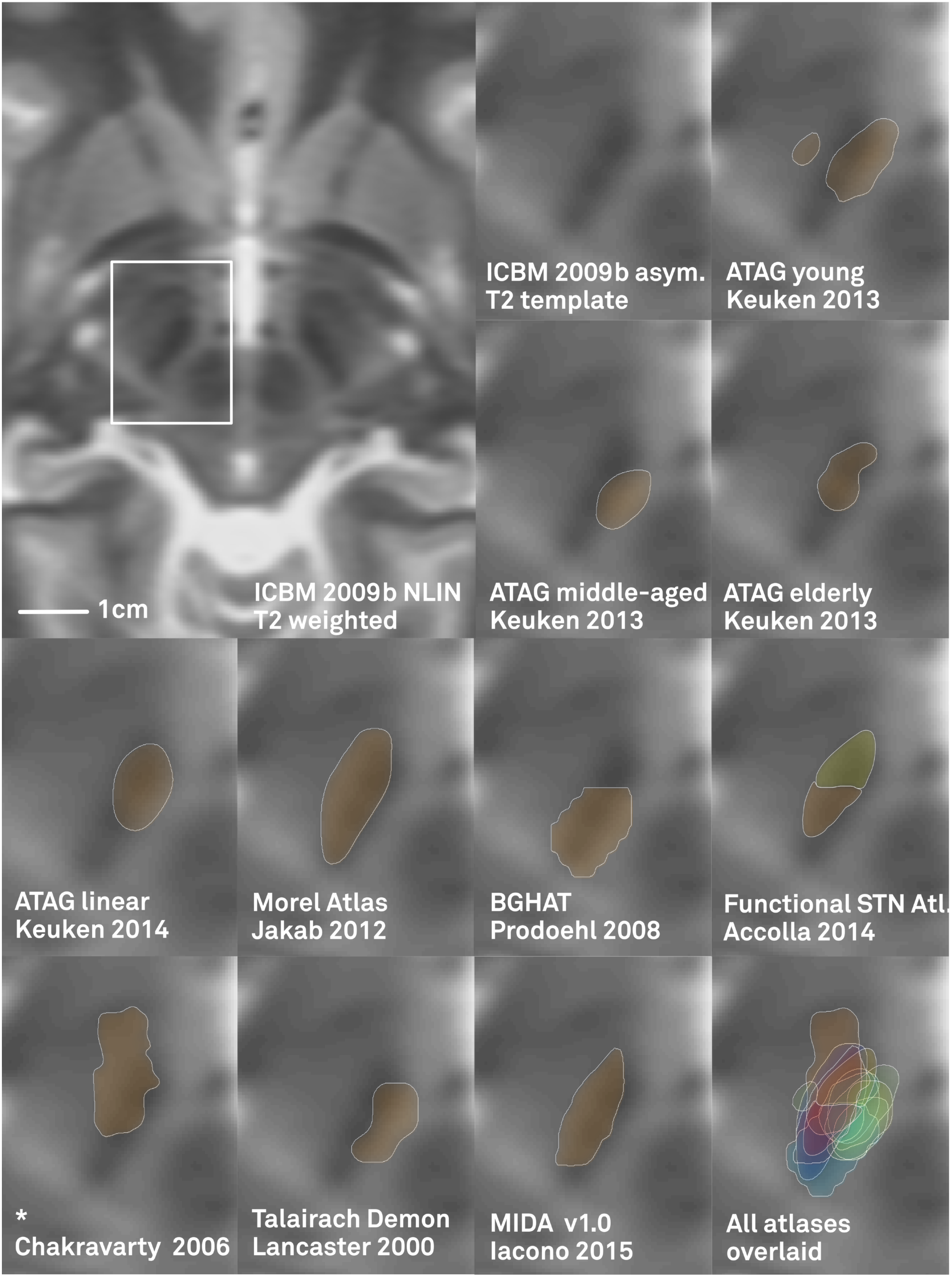
Which one is «correct»? Comparison of the ICBM 152 2009b NLIN template and various atlases of the STN available in standard space. Large panel: Overview showing selected section of the template. Small panels: First (upper middle) panel: The hypointense lentiform region of the STN defined by the template may clearly be identified. Subsequent panels show definitions of the STN based on various atlases overlaid to the template. Please note that some of the ATAG atlases have been estimated specifically to account for, e.g. age-related variance of the STN and are thus prone to vary from template anatomy (which is based on young subjects). *Also note that the Chakravarty 2006 histological atlas was coregistered to the colin27 average T1 brain template (on which the STN is not visible; Holmes et al. 1998) but is overlaid to a different template here. Even though both templates represent similar versions of the MNI space, inaccuracies on subcortical structures are prone to happen. STN definition from the Talairach Demon as defined by WFU Pickatlas (Lancaster et al. 2000; Maldjian et al. 2003). The version of the MIDA model was nonlinearly coregistered to standard space using a multimodal SyN deformation with Advanced Normalization Tools as implemented in Lead-DBS. The selected axial slice cuts through two of three functional zones of the atlas by Accolla and colleagues (motor: orange, associative: yellow). Bottom right panel shows all atlases overlaid to visualize their distribution. Axial sections displayed at z = -8 mm.

To overcome the spatial discrepancy between templates and atlases, we present a different approach to define subcortical target regions in standard space. Here, the goal was to manually define an atlas that would maximally agree with the spatial position, shape and size of the structures defined directly by the template. In order to maximize accuracy, we devised an algorithm that simultaneously used the T1- and T2-weighted as well as proton density (PD) and T2 relaxometry (T2rlx) modalities of the ICBM 2009 nonlinear templates. The algorithm created multimodal intensity fingerprints for each structure based on a few manually defined sample points and then exported probability maps that showed tissue probability for each structure in template space. These maps were then used as basis for the subsequent manual segmentation of the target regions – that was simultaneously guided by four MRI modalities.

The resulting atlas exhibits high spatial agreement to the MNI template but lacks anatomical detail and only includes structures visible on MRI. To overcome this limitation, in a second step, we used the template-based atlas as an anchor point to co-register detailed histological stacks into standard space. This has previously been done by intensity matching histologically defined labels to corresponding structures on the *colin27* MNI template (Chakravarty et al. 2008; Chakravarty et al. 2006). Using this approach, *pseudo-MRIs* defined by histology can be created and nonlinearly co-registered to the MNI templates. To increase precision of the warps especially at and around DBS regions of interest, we extended the approach introduced by Chakravarty and colleagues to include a nonlinear warp of exclusively the histologically defined DBS structures on the *pseudo-MRIs* to our template-based atlas. The two datasets, i.e. the manually segmented template-based atlas and histological stacks that were warped onto the template using the manual segmentations as additional anchorpoints were then combined to a joint dataset. The result was a patchwork atlas defining regions visible on MRI using information of the template itself but instead relying on histology for regions not discernable on MRI.

In a final step, the STN was further subdivided into three functional zones. Based on diffusion-weighted MRI acquired in ninety patients suffering from Parkinson’s disease, a tripartite segmentation of the STN into sensorimotor, associative and limbic functional zones was estimated directly within MNI space. Our final atlas constitutes a composite dataset that uses histology, structural MRI and connectivity where appropriate. The atlas both maximally agrees with the 2009b MNI space but still exhibits great anatomical detail.

## Methods

### Data acquisition

The *2009a* and *b nonlinear* versions of the *MNI 152* template were obtained from the Montreal Neurological Institute at McGill University (http://www.bic.mni.mcgill.ca/Services-Atlases/ICBM152NLin2009). The algorithm described below simultaneously worked on image acquisitions of different modalities. T1- and T2-weighted as well as proton density (PD) versions of the high-resolution ICBM 152 2009b asymmetric template and T2 relax-ometry (T2rlx) series of the 2009a symmetric template were used. A number of 20 point fiducials (*P*) were manually marked on the STN and red nucleus (RN) using 3DSlicer software (http://www.slicer.org) within MNI space. 35 points were mapped for the pallidum.

### Subcortical probability maps

The algorithm that was developed to create the probability maps as basis for subsequent manual subcortical segmentation operated on two levels. The goal of the first level was to maximize the number of tissue samples based on a low number of fiducial points *P* within the structure that were defined manually (see above). The intensity distribution of all fiducial points was sampled from each imaging modality. These values were averaged across fiducial points to form a reference intensity distribution *W* and the Euclidean distance *D* to the corresponding intensity vectors (across modalities) of each neighboring point of the starting fiducial points was calculated. Before calculating *D*, intensity values in each modality were z-scored so that distances in each modality had the same impact on *D*. The similarity *S* of each neighboring data point was calculated as follows:

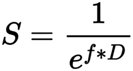

where *f* is a penalty factor that determines the harshness of the relationship between *S* and *D*. On the first level, the goal was to find as many data samples of the anatomical structure as possible. However, it would have been fatal for consecutive steps to grow the region outside the anatomical structure. Therefore, a conservative value of *f* =0.05 was empirically chosen on the first level.

Values of *S* for each point were assigned to each neighboring datapoint whereas the initial points received a score of 1. In the first run, all points that received a score higher than 0.9 were added to the set of data points *P*. This threshold was updated to amount to the mean minus one standard deviation of all points in *P* until one thousand points were assigned. From then on, the threshold was held at this fixed value. The procedure was repeated fifty times or until no additional above-threshold point could be identified. From the second iteration on, *W* was calculated by using the mean across data points *P* weighted by their scores *S*.

The second level of the algorithm worked similarly but was not based on a region growing approach like the first level. Instead, the Mahalanobis distance between intensity profiles of each voxel in the brain and the intensity covariance structure defined on the final first level point set *P* was calculated. Here, only entries of *P* that had received a score *S* above the mean plus two standard deviations of all scores were taken into account. Mahalanobis distances obtained in this way were again transferred to score values based on Equation 1 – with D now referring to Mahalanobis distance instead of Euclidean distance. Also, to include more voxels into the resulting probabilistic map, a more liberal penalty factor of *f* =0.01 was chosen.

In this part of the algorithm, a predictive value *V* was calculated for each anatomical structure and each MRI modality. This value consisted of two components. First, the ratio between standard-deviation and mean value of intensities within *P* was calculated for each MR modality (*V* _1_). The intuition of this value estimates how consistent the within-structure intensities of a certain acquisition were. For instance, if the values of pallidal points in a T2-weighted image have a low standard deviation (relative to their mean value), the within-pallidal intensities are comparably consistent. The second part, *V* _2_, instead focused on the specificity of a certain value in comparison to the rest of the brain. To assess this, the Mahalanobis distance between the mean intensity of *P* within a structure and all other points of the brain was calculated. This second score gives insight into how *rare* a certain intensity range is for a given modality, i.e. how high the classification accuracy is for a certain structure. *V* was computed by multiplying *V* _1_ and *V* _2_. *V*, *V* _1_ and *V* _2_ were normalized to add up to a sum of 1 across acquisitions (Table 1). The final *V* score gives a rough numerical estimate about how well a structure can be discerned on each sequence.

**Table 1:** Behavior of different MRI modalities in respect to the classification algorithm. First-level voxels (isotropic voxel size of 0.22 mm) describe the area covered by the reion-growing algorithm used to calculate the multimodal intensity distribution. These voxels served to form a distribution of multimodal intensities which constituted a reference for the Mahalanobis distance of each voxel on the second level. Consistency (V1), specificity (V2) and overall predictive value (V) are given for each MR sequence and each structure.

| Modality | T1 | T2 | PD | T2rlx |
| --- | --- | --- | --- | --- |
| **STN** | (711 1st level voxels included) |  |  |  |
| Mean intensities (I) | 70.36<br>±0.33 | 39.01<br>±0.44 | 56.90<br>±0.34 | 0.08<br>±0.00 |
| Consistency (V1) | 0.15 | 0.38 | 0.20 | 0.28 |
| Specificity (V2) | 0.06 | 0.49 | 0.40 | 0.05 |
| Overall predictive value (V) | 0.04 | 0.64 | 0.28 | 0.05 |
| **Pallidum** | (4699 1st level voxels included) |  |  |  |
| Mean intensities (I) | 68.12<br>±0.53 | 40.62<br>±0.42 | 61.58<br>±0.48 | 0.08<br>±0.00 |
| Consistency (V1) | 0.33 | 0.25 | 0.25 | 0.17 |
| Specificity (V2) | 0.05 | 0.65 | 0.21 | 0.09 |
| Overall predictive value (V) | 0.73 | 0.05 | 0.18 | 0.05 |
| **RN** | *(1203 st level voxels included)* |  |  |  |
| Mean intensities (I) | 68.45<br>±0.22 | 42.80<br>±0.41 | 58.82<br>±0.24 | 0.09<br>±0.00 |
| Consistency (V1) | 0.12 | 0.36 | 0.37 | 0.16 |
| Specificity (V2) | 0.06 | 0.49 | 0.40 | 0.06 |
| Overall predictive value (V) | 0.03 | 0.66 | 0.08 | 0.24 |

### Manual labelling of target structures

The manual segmentations were performed on high definition (0.22 mm isotropic voxel size) versions of the template that had been generated using 7th degree Whittaker-Shannon (sinc) interpolations as implemented in SPM12. Atlas structures were first segmented on axial slices of the template using 3DSlicer software (www.slicer.org). Boundaries of resulting labels were subsequently corrected in coronal and sagittal views. The tissue probability maps (see above) served as basis for subsequent manual segmentations. Although the algorithm that computed the tissue probability maps was not able to segment GPe from GPi completely, it defined clear borders of anatomical structures including the internal medullary lamina (Fig. 2). Given the importance of a sub-segmentation of the pallidum in deep brain stimulation, this was particularly helpful when manually segmenting these structures. For each nucleus surrounding structures were identified and the correct position of the segmented mask was verified. If necessary, it was corrected on coronal and sagittal slices. Since the STN can best be visualized on T2-weighted acquisitions as reported by several groups (Kitajima et al. 2008; Slavin et al. 2006; Dormont et al. 2004), the T2-weighted template was used as secondary guidance (together with probability maps) for segmenting STN and RN. An anatomical delineation protocol was largely adopted and modified from Pelzer and colleagues (Pelzer et al. 2013). Explicitly, borders of the biconvex lens-shaped STN were identified by localizing surrounding structures on the axial and coronal plane beginning with the red nucleus being a prominent structure on the T2-weighted template which was located medio-ventrally and in posterior direction of the STN. The anterior superior border of the STN was formed by the internal capsule. The posterior limb of the internal capsule located between GPi and STN was also marked as lateral border of the STN. The inferior posterior border was formed by the substantia nigra into which the STN was embedded and which also constituted the inferior and inferior anterior border of the STN at the level of the optic tract. The superior posterior border was marked by the zona incerta which appeared hyperintense on the T2-weighted template. The most superior part of the STN was confined by the fields of Forel (Massey et al. 2012).

**Figure 2:**
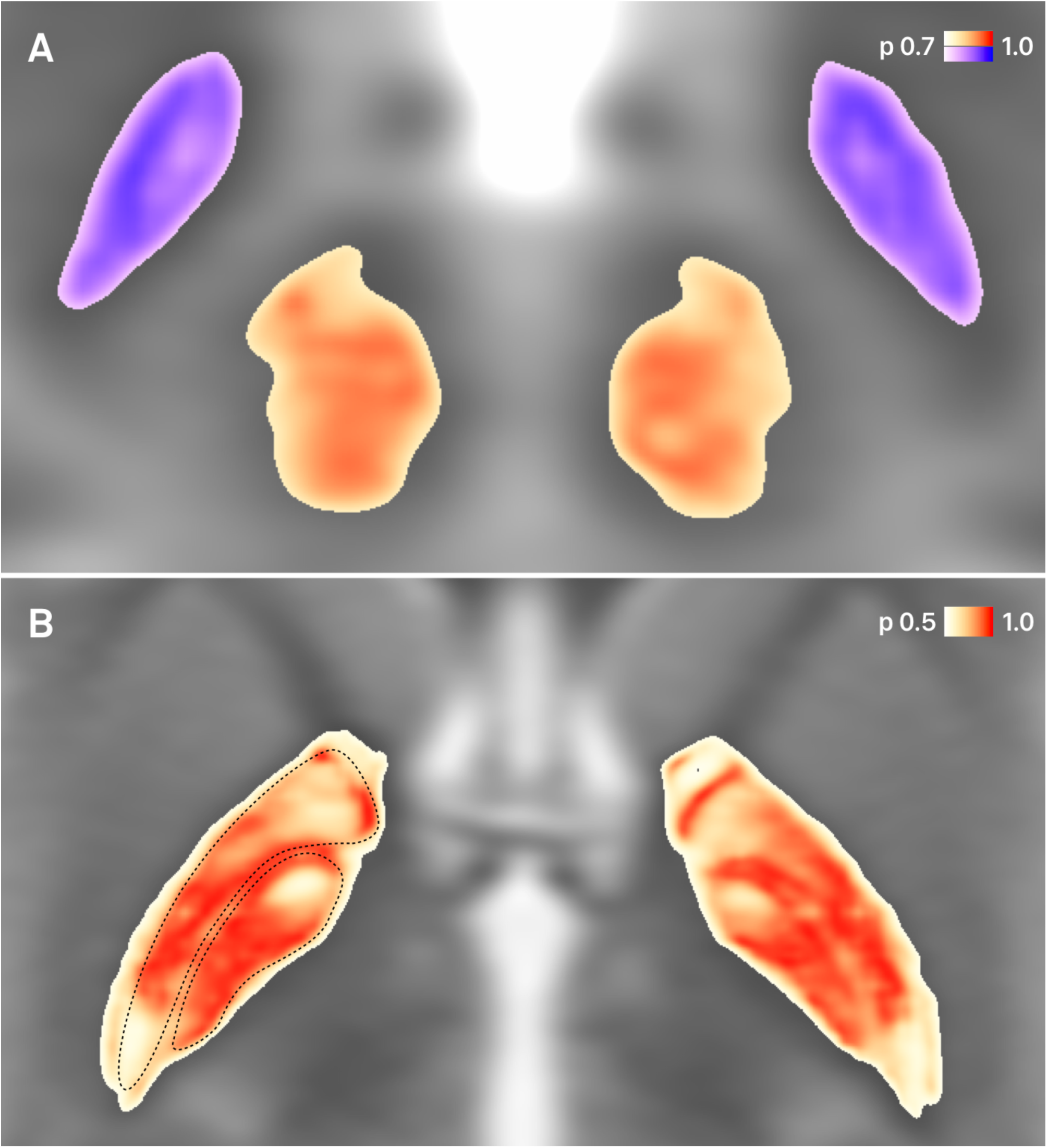
Tissue probability maps as estimated using the four modalities of the ICBM 152 template. Based on manually defined fiducial points that served as starting coordinates for a region growing algorithm in each structure, a large number of tissue sample points of the STN, RN and pallidum were generated automatically. These served as a multimodal probability distribution to which Mahalanobis distance was calculated for each voxel in the brain. A) Results for STN (cold tint) and RN (warm tint) are shown. B) Results for the pallidum are shown. Please note that the external and internal parts of the pallidum can be clearly outlined based on this computation by visual inspection. An automated segmentation between the subparts of the pallidum was not feasible given their very similar intensity distribution across acquisitions. However, based on the computational results, the two structures were segmented manually (dashed lines, shown for the right hemisphere as an example) for the final atlas. Axial slice at z = -2.6 mm.

In case of the internal and external pallidum the T1-weighted version of the template was used as secondary guidance to segment on. This decision was informed by results in predictive values *V* that ranked highest in the T1-weighted template for the pallidum (see above and results section). Again, the tissue probability maps were used as primary guidance to perform a multi modally informed manual segmentation. The GPe was separated from the GPi by the medial medullary lamina, whereas the lateral medullary lamina marked the lateral border of the GPe to the putamen. On T1, both laminae appeared more hyperintense than the pallidum. On the coronal plane, the anterior limb of the internal capsule marked the anterior-medial borders of the pallidum. Further posterior, they were partly marked by the lenticular fascicles and by the genu and posterior limb of the internal capsule. Ventrally, the pallidum was limited by the substantia innominata, the nucleus basalis Meynert, the ansa lenticularis as well as partly by the anterior commissure. Anatomical borders were in close accordance to detailed neuroanatomical atlases (Ding et al. 2016; Naidich et al. 2009; Mai et al. 2007; also Massey et al. 2012), but please note that the MNI template is not detailed enough to precisely segment especially the very fine structures that compose the surrounding structures of the targets presented here. All structures, especially GPe and GPi, were defined on individual labels without informing the process by formerly labeled other structures. Only after completing the segmentation process all four structures were projected onto the same template to assess their position relatively to each other. Due to the close spatial relationship of GPe and GPi, this functioned as an additional control to assess whether their borders toward the internal medullary lamina were symmetrically positioned. The exact definition of these borders is crucial to localize placement of DBS-electrodes in the pallidum. Slight mistakes of <1mm would have been revealed by either an overlap between the labels of GPe and GPi or by too wide a gap between their inner borders.

### Warping histology to MNI space using the template-based atlas as anchor point

The following section illustrates how the resulting template-based atlas was used as an additional anchor point to coregister a histological atlas to ICBM 2009b NLIN space. Given that the template-based atlas consisted of core target regions of the DBS context, it can especially contribute to a high warping precision of these targets. Here, we compared two multimodal warp estimates using the Advanced Normalization Tools (ANTs; http://stna-va.github.io/ANTs/), one including and one excluding the template-based atlas as anchor point. The subcortical atlas developed by Chakravarty and colleagues was originally derived from a single set of high-resolution, thin-slice histological data of the region of the basal ganglia and thalamus in a single brain (Chakravarty et al., 2006). It contains 90 anatomical structures that were manually delineated by an experienced neuroanatomist. The histological and geometric data was reconstructed in 3D, slice-to-slice coregistered and manually intensity matched to the T1 and T2 templates, forming *pseudo-MRI* images in T1 and T2 weighting. These coregistered *pseudo-MRI* images formed the basis for the first multi modal normalization to the 2009b NLIN space using ANTs (ω_1_). In a second approach, the regions defined by the template-based atlas were included into the nonlinear deformation as a third transformation stage. This transform is referred to as ω_2_ in the following (see Fig. 3).

**Figure 3:**
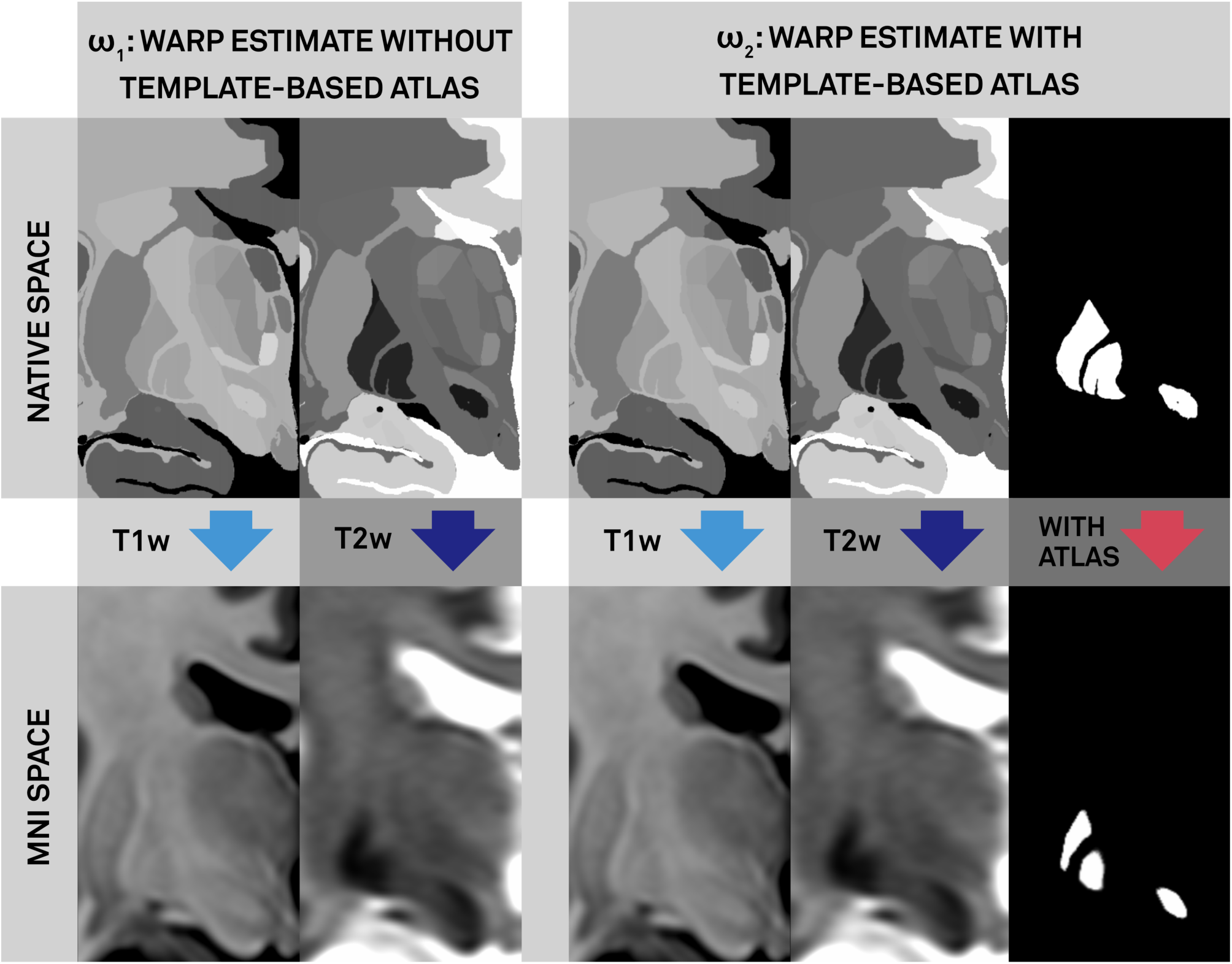
The warping mechanism using the Advanced Normalization Tools (ANTs http://stnava.github.io/ANTs/) in a multi modal approach is displayed. ω1: Both the T1-weighted and T2-weighted pseudo-MRI images based on the histological atlas digitalized by Chakravarty et al. (2006) are used to create a final warp estimate for the normalization of the original atlas to MNI space (ICBM 2009b NLIN). ω2 displays the second approach using the template-based atlas as an additional anchor point when estimating the warp: A binarized image of the original histological Chakravarty atlas displaying solely the labels corresponding to the structures included in the template-based atlas (GPe, GPi, STN, RN). ANTs estimates the warp field of this source image onto the template-based atlas in MNI space and adds this information to the warp estimate from the T1- and T2-warp estimate to compute a joint multi-modal deformation field.

### Segmentation of STN based on diffusion-weighted MRI

Structural and diffusion-weighted acquisitions of 90 PD patients were downloaded from the Parkinson’s progression markers initiative (PPMI) database PPMI database (mean age 61.38 ± 10.42 SD, 28 female). T2-weighted images had an in plane resolution of 0.94 × 0.94 and a slice thickness of 3 mm. DWI acquisitions had an in plane resolution of 1.98 × 1.98 and a slice thickness of 2 mm and were acquired with following parameters: TE/TR 61/900 ms, 64 gradient directions, flip angle: 90°, field strength: 3T. Detailed scanning parameters can be found on the project website (www.ppmi-info.org). For each subject, a nonlinear deformation field into ICBM 2009b NLIN asymmetric space was estimated based on T2-weighted acquisitions using a fast diffeomorphic image registration algorithm (DAR-TEL) as implemented in SPM12 (Ashburner et al., 2007; http://www.fil.ion.ucl.ac.uk/spm/software/spm12/) and Lead-DBS (Horn & Kühn 2015; www.lead-dbs.org). Global fibersets were calculated using a generalized q-sampling method as implemented in DSI-Studio / Lead-DBS (Yeh et al., 2010; http://dsi-studio.labsolver.org/). For each patient, 20k fibers were sampled seeding within a white-matter mask that was defined by segmenting the T2-weighted acquisitions using SPM12. For each patient, the fiberset was then normalized into MNI space following the approach described in (Horn et. al, 2014; Horn & Blankenburg 2016). The segmentation pipeline used to segment the STN functional zones is described in (Horn & Blankenburg 2016). Briefly, for each voxel of the STN, connectivity strength to motor, associative and limbic regions was calculated and voxels were assigned to either functional zone using a winner-takes-all approach. Motor, associative and limbic cortical regions used as connectivity seeds were largely informed by the study of Accolla and colleagues (Accolla et al. 2014) but were based on multiple whole-brain atlases (see S2 for details).

## Results

The algorithm robustly produced useful tissue probability maps for each of the four structures. It first started based on a low number of manually defined sample points and grew a region around these points based on multi modal tissue intensities. This was done solely to enlarge the sample size of voxels surely residing in each structure. The final numbers of voxels included in this region growing algorithm are denoted in Table 1. Mean intensity values used as *P* on the second level are denoted as *I.* These values formed the distribution to which Mahalanobis distances for each brain voxel were computed to. Additionally, consistency and specificity values (*V* _1/2_) as well as overall predictive values (*V*) were computed for each target and sequence. The MNI template showing the best overall score to discriminate the STN was the T2-modality followed by PD. The same order applied to the RN, whereas for the pallidum, as expected, T1 followed by PD yielded highest scores. Probability maps showing probability scores based on Mahalanobis distances to region-defining intensity distributions are visualized in Fig. 2. Please note the clear differentiation between internal and external segments of the pallidum. Manual segmentations of the four structures (STN, RN, GPi, GPe) were subsequently performed on a high definition (0.22 mm isotropic voxel size) version of the template using the tissue probability maps as primary guidance. In total, 63, 43, 48 and 36 axial slices were manually labeled for GPe, GPi, RN and STN respectively. The final STN segmentation for left and right STN yields a sum of 26.272 voxels each 0.0106 mm^3^ in size resulting in a volume of approximately 140 mm^3^ per STN nucleus. GPe parcellations had a volume of approximately 929 mm^3^, GPi of 382mm^3^ and the RN of 316mm^3^ per nucleus. In some border areas of the target structures and especially when delineating the GPe from GPi, the high signal-to-noise contrast of the tissue probability maps constituted a very helpful resource in the manual labelling process (S2). As pointed out by Jovicich and colleagues, the accurate and reliable measurement of subcortical brain volumes from MRI data is a nontrivial task (Jovicich et al. 2009). Moreover, due to the high resolution of the final output, the process amounted to ~120 hours of concentrated manual labelling. Despite this substantial effort, a high resolution of template space was chosen to attain high final resolution and more robustness due to redundant relabelling on each of two adjacent slices. Final results of the manually segmented atlas are visualized in Fig. 4.

**Figure 4:**
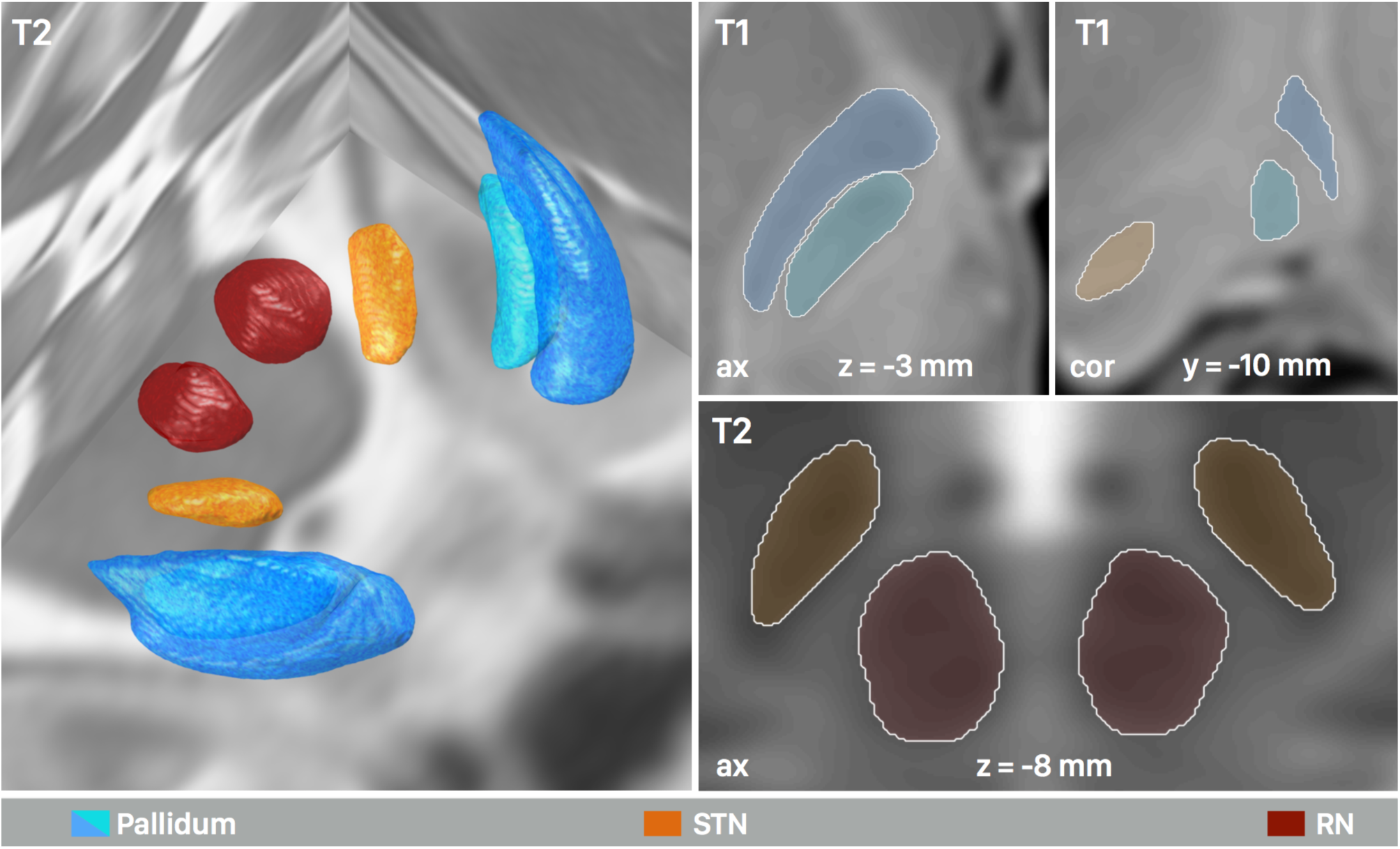
Final segmentation of the template in 3D (left half) and 2D (right half) as informed by the automated probability maps to maximize objectivity. Here, the atlas is shown as implemented into the Lead-DBS toolbox (www.lead-dbs.org). Structures were defined in a 0.22 mm isotropic resolution.

To fill in the gaps of the template-based atlas with further anatomical detail, a histology based and digitized histological atlas (Chakravarty et al., 2006) was co-registered into standard space. This was done using two methods, i) the method originally introduced by Chakravarty and colleagues that generates *pseudo-MRIs* based on histology which are nonlinearly warped onto the MNI templates (ω_1_) and ii) the same method that was extended by our template-based atlas as an additional anchor point (ω_2_) (Fig. 3). The overlap of each warped nucleus with its ground truth (defined by manual segmentations) was assessed computing Cohen’s Kappa κ (Cohen, 1960; McHugh, 2012). For the pallidum, κ(ω_1_) was 0.67 (interpreted by Cohen as substantial agreement) vs. κ(ω_2_) was 0.82 (perfect agreement). Since one of the pallidum labels of the histological atlas contained parts of the GPe and GPi (label 5) it was not feasible to further discriminate between the external and internal part of the pallidum when computing Cohen’s Kappa. For the STN κ(ω_1_) was 0. 41 (moderate agreement), and κ(ω_2_) was 0.73 (substantial agreement). For the RN κ(ω_1_) was 0.69 (substantial agreement), and κ(ω _2_) was 0.92 (perfect agreement). Fig. 5 shows a direct comparison between the results of the two warp estimates which clearly illustrates the benefit of ω_2_ over ω_1_.

**Figure 5:**
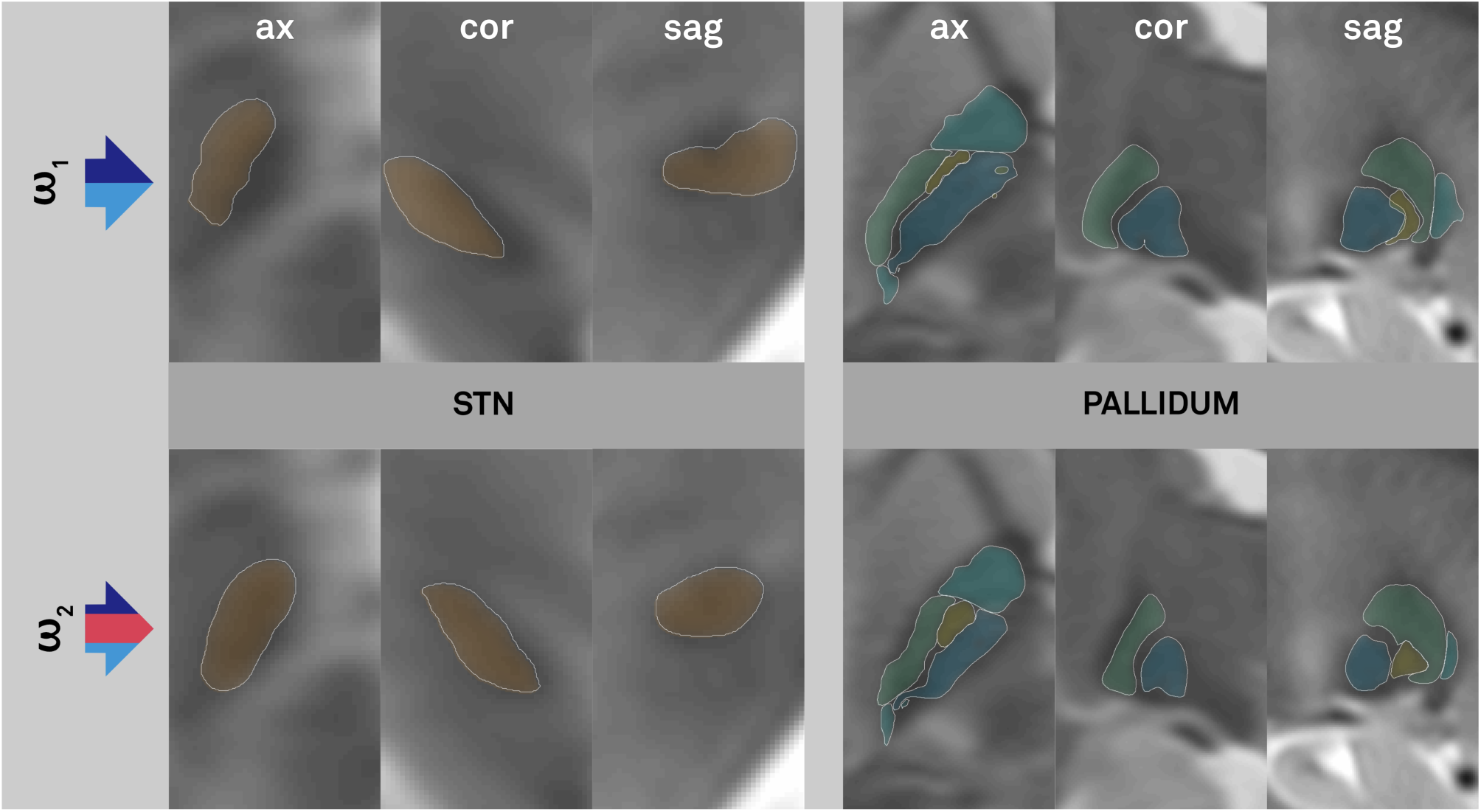
Results of the two different warp estimates ω1 and ω2. The colored arrow on the left symbolizes the different components of each warp estimate as explained in Fig. 3. The upper panel shows the results of the warped histology atlas (Chakravarty et al., 2006) when using a multimodal (T1/T2) SyN registration to nonlinearly warp the structures onto the MNI templates (extended approach similar to the one used in original study by Chakravarty and colleagues). The lower panel shows the results of the same atlas where the method was further extended by including the template-based atlas as an additional anchor point to estimate the warp field.

Sub-segmentation results based on fiber tracking from cortical functional zones (see S2 for details) to the STN resulted in a tripartite segmentation of the STN comparable to results from prior studies (Fig. 6; Accolla et al., 2014, Lambert et al., 2012). Namely, the dorso-lateroposterior region of the STN was assigned its sensorimotor functional zone, the dorsal mid portion the associative zone and the ventroanteriormost portion the limbic zone. Such a parcellation is well-known from the literature but adds further anatomical detail to the MRI-/histology-/tractography composite atlas presented in this study. Fig. 6 summarizes the fiber tracking results. Also, here, it was performed directly within MNI space following the approach validated in (Horn et al., 2016).

**Figure 6:**
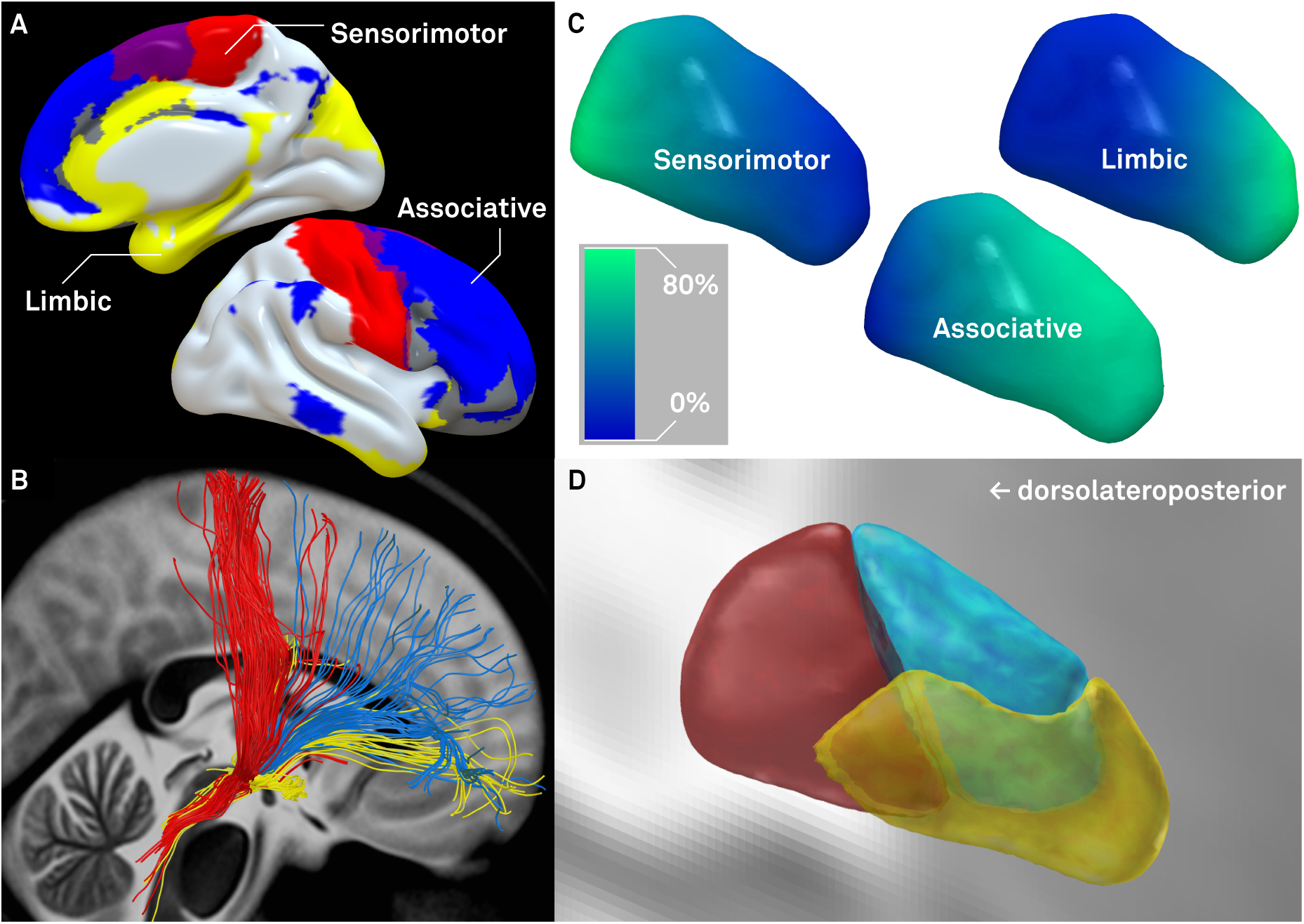
A) Cortical regions used for fibertracking (see S2 for details) – magenta shows overlap between sensorimotor and associative seeds, grey between associative and limbic seeds, B) Fibers selected from the group connectome based on 90 PD patients that exclusively connect cortical seeds to the STN, C) Percent fibers received by STN voxels originating from the three functional zones mapped to the surface of the STN, D) Final winner takes all parcellation of the STN.

On each hemisphere, the final composite atlas consisted of 92 regions in total. Of these 92 regions three were based on structural connectivity and the MNI template segmentation, namely the tripartite STN. Another three regions were solely based on the MNI template segmentation, namely GPe, GPi and RN. Finally, the remaining 86 regions were defined by histology atlas. Fig. 7 summarizes the three-layered construction of the composite atlas by means of a step-by-step flowchart (A-D) and visualizes the final atlas in two-dimensional sections (E).

**Figure 7:**
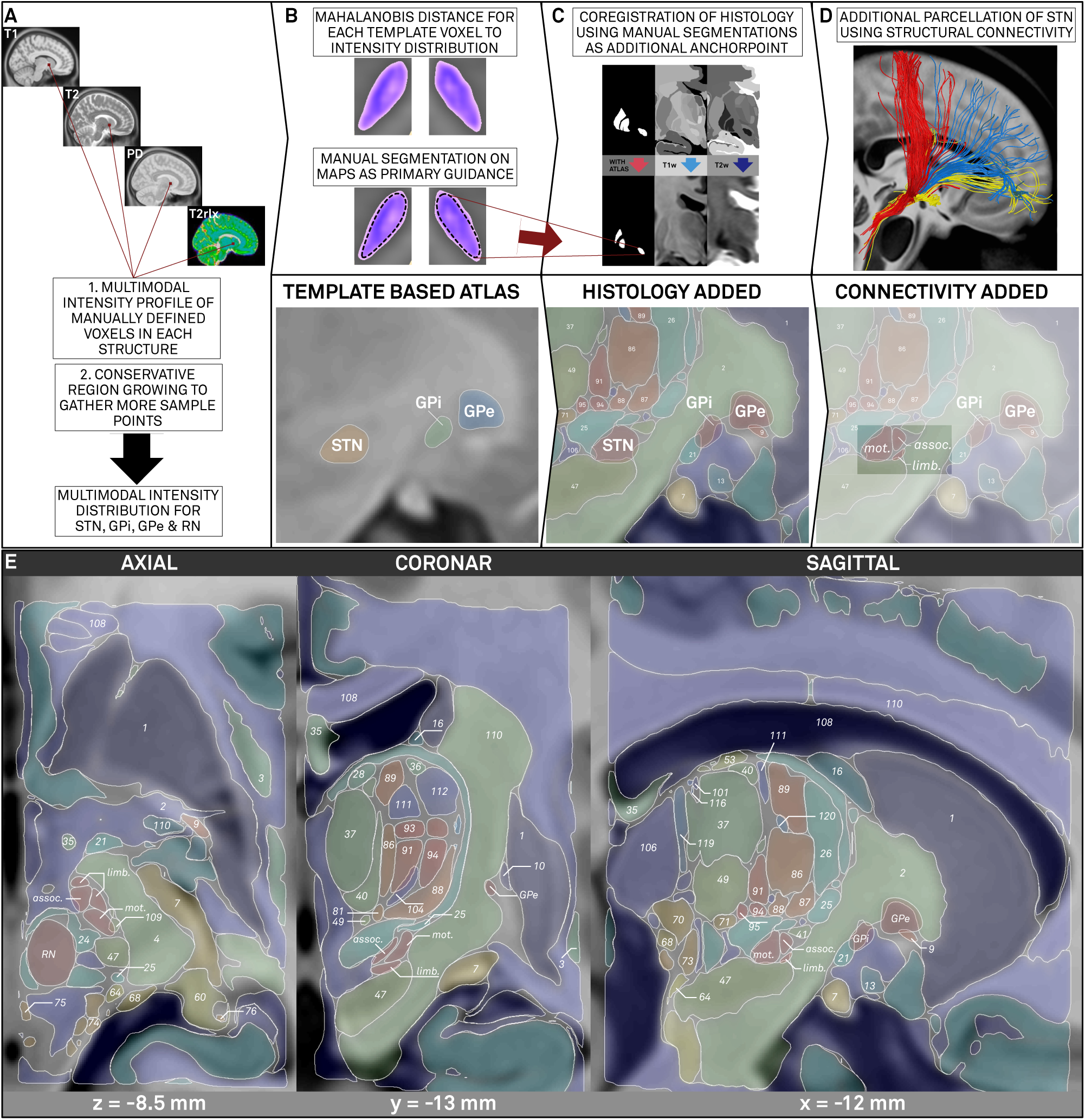
Overview of the construction and results as displayed on the MNI template of the composite atlas. A-D) Construction of the three-layered composite atlas. A) Multimodal tissue probability maps are computed. B) These maps served as primary guidance for the manual segmentation of the target structures on the MNI ICBM 2009b NLIN template which constitute the first component of the atlas. C) As a second component a histological atlas was added and normalized to the MNI space using the formerly segmented target regions as additional anchor point to complement the warp estimate of remaining atlas structures. D) A third component was added by the functional segmentation of the STN based on tractography into motor (mot.), associative (assoc.) and limbic (limb.) functional zones. E) Results of the final composite atlas consisting of 92 labels in total* with segmented STN, Regions manually segmented from the MNI template series (RN: red nucleus, GPi/GPe: internal/external parts of the pallidum) as well as remaining anatomical structures defined by histology (denoted by numbers; for label references see Chakravarty et al., 2006). (*Not all numbers have been issued)

## Discussion

Three main conclusions may be drawn from this study. First, we developed an algorithm that combines images from different MR modalities into a single probability map that is helpful in manual segmentation processes. By using these probability map in a subsequent manual segmentation step, two prominent DBS targets were segmented directly in MNI space, using the integrated information of four MR-modalities (T1, T2, proton density and T2-relaxometry). Second, we demonstrated a first application of such an atlas by using it to accurately deform a more detailed, histology-based atlas into MNI space. The resulting patchwork-atlas uses the template to define regions that can be outlined on MRI alone and uses additional histology to define remaining regions. Finally, we added a third component to the atlas that segments the subthalamic nucleus into three functional zones based on their structural connectivity to cortical regions. The final “MRI-histo-connectomic” atlas constitutes a detailed description of the subcortex that uses the most accurate source of information for each region and is in perfect alignment with the ICBM 2009b nonlinear asymmetric template.

As a first component of the atlas, primary targets for deep brain stimulation, the STN and pallidum, as well as the red nucleus were segmented on the template, given the great clinical interest in these structures. The latter was additionally selected because a high robustness of the spatial relationship between RN and STN has been described and because the RN is conventionally used as a waypoint in surgical planning for STN-DBS (Andrade-Souza et al. 2008; Starr et al. 2002; Pollo et al. 2003 but also see Danish et al. 2006). Several subcortical atlases exist that stem from various sources (Prodoehl et al. 2008; Keuken et al. 2013; Qiu et al. 2010; Jakab et al. 2012; Chakravarty et al. 2006; Yelnik et al. 2007; Krauth et al. 2010; Keuken et al. 2014; Morel 2013). The anatomical detail of some of the available atlases goes far beyond the segmentations feasible directly on the MNI template. Still, a proper spatial definition of DBS targets *within* the standard template was a crucial step to perform co-registrations of histology to MNI space. Also, since the template is used for nonlinear warps in DBS imaging, a proper definition of DBS targets on the template itself is a prerequisite for group studies that analyze the relationship between spatial locations of DBS electrodes with respect to anatomical targets and clinical outcomes. In comparison to single subject analyses, these studies depend on spatial normalization into common space and a proper and observer-independent definition of the anatomical target herein. Before, studies with similar goals have overcome the issue by nonlinearly coregistering patient data to the space of a certain atlas (instead of the ICBM template; e.g. Yelnik et al. 2007; Welter et al. 2014; Butson et al. 2007) or by segmenting the STN into geometrical subfields within each patient’s native space (Herzog et al. 2004; Wodarg et al. 2012). A limitation of these approaches is that they do not use the full potential of years in research that went both into developing standardized, study-independent and well-characterized anatomical templates (Fonov et al. 2011b; Fonov et al. 2009; Allen et al. 2002) and into the steadily ongoing evolution of nonlinear deformation algorithms (Ashburner 2012; Ashburner 2007; Avants et al. 2011; Avants et al. 2008; Klein et al. 2009). Even more importantly, results are not generalizable to novel patient populations if the atlases and/or atlas templates are not broadly available.

Furthermore we could show, that the atlas presented here can assist in the process of warping more detailed atlases into standard space. More specifically, datasets defined on histological stacks (Morel 2013; Yelnik et al. 2007; Mai et al. 2007), like the one used to test our hypothesis (Chakravarty et al. 2006), could be coregistered into MNI space using additional spatial information of our atlas as anchor points. Histological atlases give detailed insight into surrounding structures of DBS targets that played a prominent role in DBS targeting research, such as the zona incerta (Karlsson et al. 2011; Schmitz-Hubsch et al. 2014; Plaha et al. 2006; Blomstedt et al. 2012) or the dentatorubrothalamic tract (Meola et al. 2015; Coenen et al. 2011). But if their definition of the main targets does not exactly match the ones of the template used for nonlinear coregistration, the information gained from the use of such atlases may be misleading. In their original study, Chakravarty and colleagues introduced a novel method that generates *pseudo-MRIs* based on histological labels that could then be nonlinearly co-registered to a template. Here, we extended this method by an additional third warp that included our template-based atlas. As can be seen in Fig. 5, results of this extended approach define the areas of main focus (i.e. the DBS targets) with much greater accuracy and fill the gaps not defined by the template-based atlas with information derived from histology.

Finally, the subthalamic nucleus was further parcellated using structural connectivity. This has been done before (Accolla et al., 2014; Lambert et al., 2012) and cortical seed definitions used in the present study are largely informed by the study of Accolla et al. However, in comparison to their study, we added several improvements to the method. First, fiber tracking information was derived from a population of 90 PD patients (vs. 13 in Accolla et al.). Second, we applied a nonlinear normalization approach to warp fiber tracts into MNI space that showed greater accuracy in a head-to-head comparison (Klein et al., 2009) and was validated to warp fiber tracts into standard space (Horn et al., 2016).

Our final atlas may be used to visualize spatial positions of DBS electrodes in context of subcortical anatomy after reconstruction using e.g. Lead-DBS software. The addition of histology to the purely template-based segmentation allowed us to include various other DBS targets such as the ventral intermediate nucleus (VIM) commonly used to treat essential tremor. Fig. 8 shows an example visualization of three common DBS trajectories using a selection of atlas structures relevant to the field of DBS. Such a visualization may be used for postoperative programming of DBS devices. Please note, that further additional DBS targets such as the nucleus ventro-oralis intermedius (Tourette’s syndrome) or caudal zona incerta (alternative target in PD) and multiple surrounding structures associated with side-effects (e.g. internal capsule, optic tract, GPe) are also defined by the atlas.

**Figure 8:**
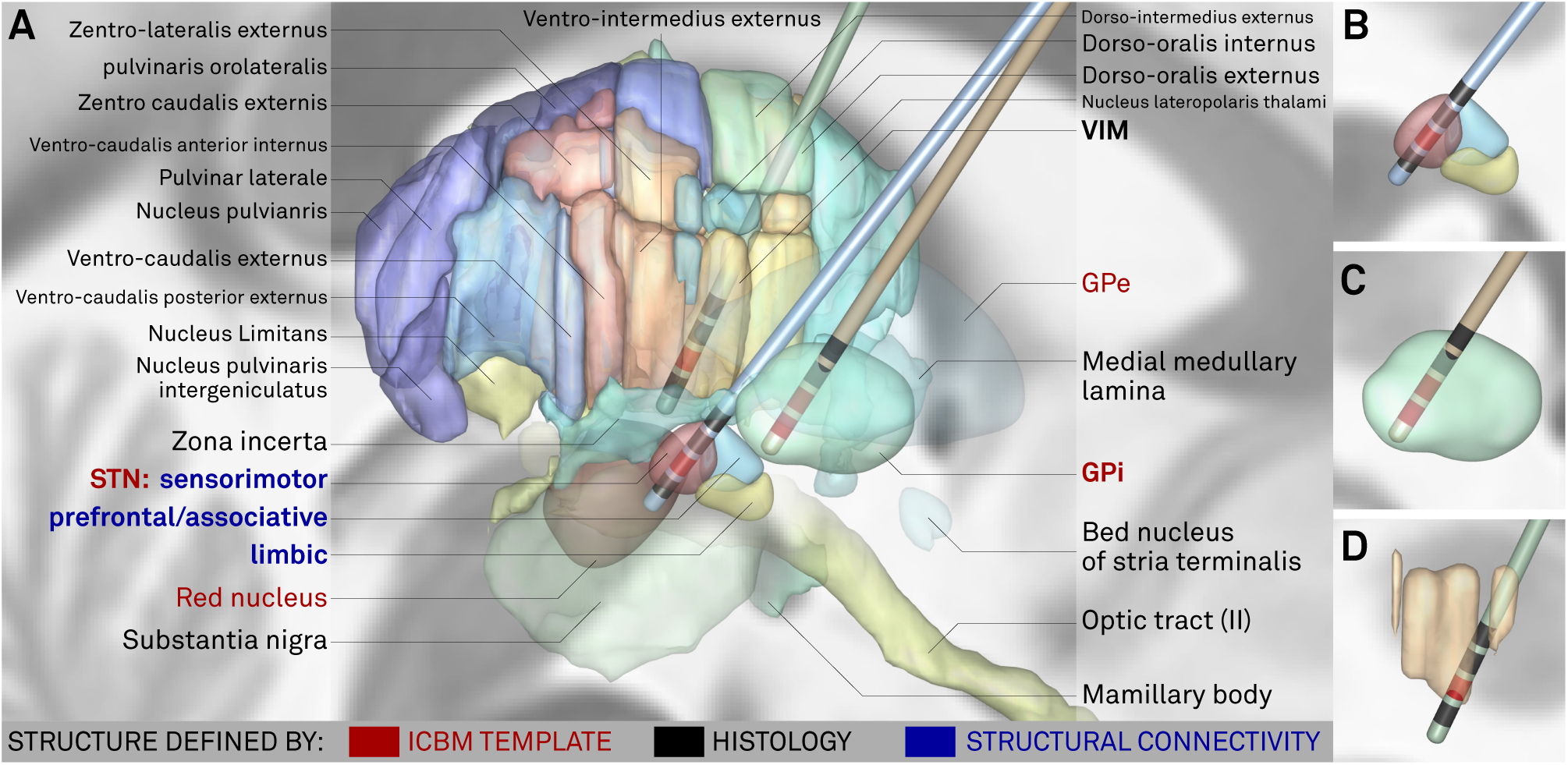
Visualization of three typical DBS trajectories in synopsis with structures of final composite atlas relevant to DBS. Based on spatial relationship to atlas structures, postoperative programming could be performed, e.g. selecting contacts highlighted in red as active cathodal contacts. DBS target structures marked in bold. Please note that the final atlas consists of numerous structures not visualized in favor of clarity, some of which with potential interest in DBS (such as e.g. internal capsule, campus Forellii). Some structures (GPe, VIM, bed nucleus, medial medullary lamina, optic tract, substantia nigra) were rendered (partly) more opaque than others to put focus on most relevant parts of the scene. overview with labels, B) close-up of STN trajectory with functional zones, C) GPi trajectory and D) VIM trajectory.

As an important limitation to our atlas, we would like to raise a different question. In the caption of Fig. 1, we asked *which* atlas correctly defined the STN. However, for many applications, this is the wrong question to ask altogether. Instead, «*what* is correct?» should be asked. A good example is the ATAG dataset of STN atlases which exists for three age populations (Keuken et al. 2013). In our view, both its publication and release under an open license were extraordinarily valuable contributions to the neuroscientific community since they clearly showed that position and shape of the STN varies with subject age and made the data available for everyone to examine directly. Thus, while our atlas tries to maximize overlap with the ICBM template, the approach of Keuken and colleagues was the opposite: to uncover and define anatomical variability of the nucleus. We argue that both goals are valid and may help to answer the questions for which they were designed. A prerequisite of the (absolute) validity of our approach is the (absolute) correctness of normalization results obtained from nonlinear deformation algorithms. Of course, despite the increasing accuracy of the latter, this prerequisite is not fulfilled in practice. Most non-linear deformation algorithms even introduce penalty equations that prevent them from extreme overfitting and high inhomogeneity of deformation fields at some point. Another reason is that DBS targets may not exhibit a large enough contrast for the algorithm to create an accurate deformation – especially when using MRI data acquired in a clinical setting. Thus, datasets like the one introduced by Keuken and colleagues may be of great value when studying a certain age population.

Another important limitation includes the partial disagreement between the STN when visualized using MRI and histology (also see S1). The dorsolateral part of the nucleus (which corresponds to its sensorimotor functional zone) does not render as hypointense as the ventromedial part (Dormont et al. 2004; de Hollander et al. 2014; Richter et al. 2004; Schäfer et al. 2011, Massey et. al 2012). Notably, the volumes of the STN in MRI studies (e.g. 31-72 mm^3^ in Schäfer et al. 2011) are substantially lower than in histological studies (e.g. 240 mm^3^ in Hardman et al. 2002; also see Richter et al. 2004 for a direct comparison). The STN volume of the present study was 140 mm^3^ and is thus closer than usual to reality. Still, this limitation is especially important in the present case since our approach used MRI intensities (on the template) to co-register histology. Thus, based on prior results, one may assume that the STN shown in the final atlas is too small, especially in the dorsolateral part. Since our aim was to construct an atlas that could be used to control DBS electrode placement, this limitation is very important. On the other hand, DBS placement controls in clinical routine often equally use MRI to detect the STN within native anatomical space of a patient and here, of course, the same limitation applies. Still, as an outlook, we plan to extend the atlas presented here based on paired histology-MRI datasets in future studies and release a revised version of the atlas subsequently.

In this study, a subcortical atlas was defined by direct manual segmentation of the ICBM 2009 152 template series in high resolution. The manual segmentation process was largely informed by automatically generated probability maps that integrated the intensity information of four template modalities (T1, T2, proton density, T2 relaxometry). Subsequently, a detailed histological atlas was co-registered to fill gaps with anatomical structures not visible on MRI. Finally, given its great importance in the DBS literature, the STN was further sub-segmented into three functional zones using structural connectivity to the cortex. At the present time, the structures defined by the template and by structural connectivity will be made publicly available to the scientific community under an open license using the name DISTAL atlas (**D**BS **I**ntrin**s**ic **T**emplate **A**t**L**as) and distributed within the software package Lead-DBS (www.lead-dbs.org).

## Acknowledgments

The study was supported by the German Research Agency (DFG - Deutsche Forschungsgemeinschaft). Grant Number: KFO 247. 

SE and AH received funding from Stiftung Charité, Max-Rubner-Preis; AH further received funding from Berlin Institute of Health and Prof. Klaus Thiemann Foundation.

Data used in the preparation of this article were obtained from the Parkinson's Progression Markers Initiative (PPMI) database (www.ppmi-info.org/data). For up-to-date information on the study, visit www.ppmi-info.org. PPMI - a public-private partnership - is funded by the Michael J. Fox Foundation for Parkinson's Research and funding partners, see www.ppmi-info.org/fundingpartners.

## Supplemental material

### Existing atlases of the STN do not exactly match the MNI template (S1)

As can be seen in Fig. 1, existing atlases do not necessarily match the spatial position, shape and size of the STN defined by the MNI template. Various reasons may explain this disagreement. First, some atlases have been registered to different brain templates than the ICBM standard (ICBM 152 2009 nonlinear template series; Fonov et al. 2011). Various spatial standard templates exist, most of which represent a function of the historical development of nonlinear warping techniques (Ashburner 2012; Fonov et al. 2011; Fonov et al. 2009; Allen et al. 2002) or of differing underlying populations (Liang et al. 2015). Even two MNI template series labeled as *MNI152* exist. The first one is referred to as the 6th-generation nonlinear template and is the successor of the *MNI305* (Collins 1994). It was adopted as the standard template for brain anatomy by popular fMRI processing suites such as SPM (http://www.fil.ion.ucl.ac.uk/spm/; since version SPM99) and FSL (http://fsl.fm-rib.ox.ac.uk/fsl/fslwiki/). Here, MRI acquisitons of 152 subjects were linearly and nonlinearly coregistered to the *MNI305* space (see Brett et al. 2002 for an overview). In the newest version to date (2009 series), the acquisitions were nonlinearly coregistered using updated algorithms (Fonov et al. 2011b). In contrast to their 6th-generation predecessor, they exhibit much greater detail and signal to noise ratio. Notably, within this current most series, the 2009b template is the only one exhibiting a resolution as high as 0.5mm. As a parallel offspring, the MNI generated a single-subject template in similar space by scanning one subject 27 times. This template is referred to as the *colin27* space and is again used to define single subject anatomy in SPM and FSL. Lastly, templates that are based on populations of different ethnicity (e.g. see Liang et al. 2015) or ones that focus on parts of the brain such as the cerebellum (Diedrichsen et al. 2011) exist. All of these standard templates roughly define the space that is often simply referred to as the *MNI space*. In the field of fMRI imaging, where spatial resolution of most studies ranges within the magnitude of 2-3 mm, slight inaccuracies introduced by a mixed use of templates may not lead to significant errors. In DBS imaging, however, millimeters do matter (Frankemolle et al. 2010; Yelnik et al. 2003; Horn & Kühn 2015b; Pollo et al. 2004), especially when studying the spatial relationship between electrode and target region. It is important to note that in the field of DBS imaging the precise warping and necessary deformation of a patient’s STN entails the same deformation of the DBS electrode placed within or near the STN. This however might result in a DBS electrode that may appear smaller or larger in the standard space as compared to the native space. In some cases, they might even form a bend (supplementary Fig. 1). In MNI space, however, the STN itself should always appear at the same position and have the same shape and size: the one of the template.

Further reasons that may explain slight disagreements between available atlases and the ICBM 2009b nonlinear template anatomy may be that coregistration algorithms led to inaccuracies of structure placement or that the number of volumes acquired for atlas generation differed from the ones acquired for template generation. For instance, one version of the ATAG atlas is based on 30 subjects with a mean age of 24.2 (Keuken et al. 2014) whereas subjects of the MNI 152 database range from birth to adulthood (Fonov et al. 2011b). Where the MNI 152 2009 nonlinear templates were constructed using nonlinear transforms estimated by MINC Tool Kit software (http://www.bic.mni.mcgill.ca/Services-Software/ServicesSoftwareMincToolKit), the ATAG atlas was constructed using linear AC/ PC transforms (which are comparable to Talairach transforms (Brett et al. 2002)). Ultimately, a different spatial definition is prone to result given the large differences in data used and methods applied and it is important to mention that this was exactly the goal of the ATAG experiment: To uncover variability of basal ganglia anatomy (and not to segment basal ganglia from the ICBM template). Lastly, some reasons for incongruence between atlas and template structures should be attributed differently and again, the STN is a good example. Given an inhomogeneous distribution of iron-density in the nucleus, its medioventral portion shows up more hypointense than its dorsolateral parts (Dormont et al. 2004; de Hollander et al. 2014; Schäfer et al. 2011; Richter et al. 2004). This may lead to a smaller representation of the STN on standard MRI and greatly justifies the use of histological atlases or imaging techniques that are more sensitive to iron, such as quantitative susceptibility mapping (Wang & Liu 2015). Nevertheless, a meticulous coregistration between atlas sources and the standard template as well as the use of the same template in atlas generation and subsequent application procedures is important.

### Definition of seeds for STN segmentation (S2)

The STN was segmented into sensorimotor, associative and limbic functional zones largely following the study of Accolla and colleagues (2014). As in their original study, sensorimotor regions included primary motor and sensory cortices as well as premotor cortices and supplemental motor area. Associative regions consisted of prefrontal cortex including superior, middle and inferior frontal gyri. Finally, limbic regions included ACC, hippocampus, amygdala and orbitofrontal cortex including medial, anterior, posterior and lateral orbitofrontal cortex. To assure maximal observer and dataset independence, corresponding regions in seven whole-brain parcellations were used to define these regions. All of the atlases are available within the Lead-DBS software package (www.lead-dbs.org). Table S1 summarizes the region indices used to define sensorimotor, associative and limbic regions for each atlas set. Seeds were weighted in terms of an N-image, i.e. voxels that were defined by all seven atlases received a score of seven, and so forth. Final maps are depicted in Fig. 6.

**Table S1:**
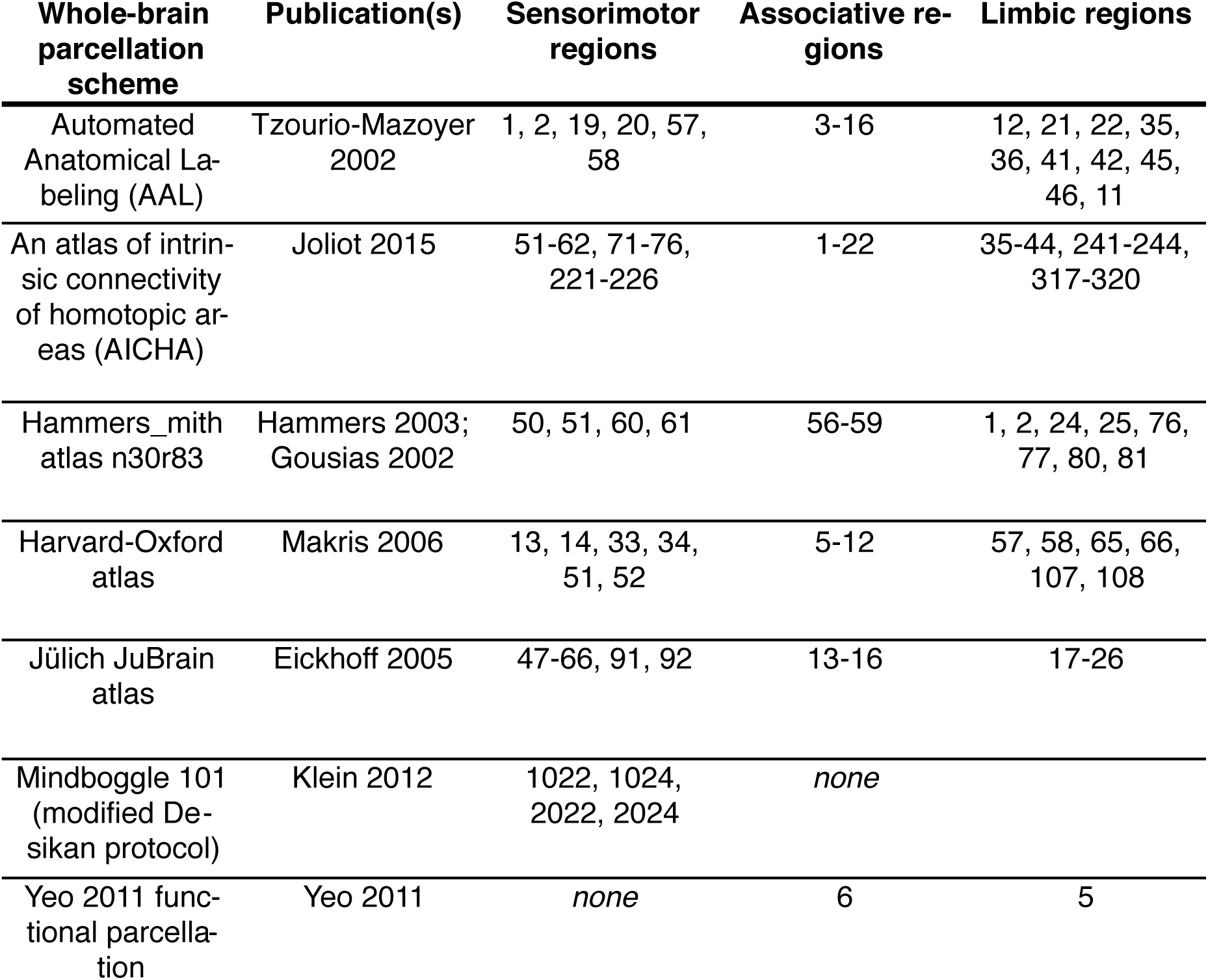
Regions of various whole-brain parcellations included to form functional seeds for STN segmentation

| Whole-brain parcellation scheme | Publication(s) | Sensorimotor regions | Associative regions | Limbic regions |
| --- | --- | --- | --- | --- |
| Automated Anatomical Labeling (AAL) | Tzourio-Mazoyer 2002 | 1, 2, 19, 20, 57, 58 | 3-16 | 12, 21, 22, 35, 36, 41, 42, 45, 46, 11 |
| An atlas of intrinsic connectivity of homotopic areas (AICHA) | Joliot 2015 | 51-62, 71-76, 221-226 | 1-22 | 35-44, 241-244, 317-320 |
| Hammers_mith atlas n30r83 | Hammers 2003; Gousias 2002 | 50, 51, 60, 61 | 56-59 | 1, 2, 24, 25, 76, 77, 80, 81 |
| Harvard-Oxford atlas | Makris 2006 | 13, 14, 33, 34, 51, 52 | 5-12 | 57, 58, 65, 66, 107, 108 |
| Jülich JuBrain atlas | Eickhoff 2005 | 47-66, 91, 92 | 13-16 | 17-26 |
| Mindboggle 101 (modified Desikan protocol) | Klein 2012 | 1022, 1024, 2022, 2024 | *none* |  |
| Yeo 2011 functional parcellation | Yeo 2011 | *none* | 6 | 5 |

### Supplementary figures

**Figure S1:**
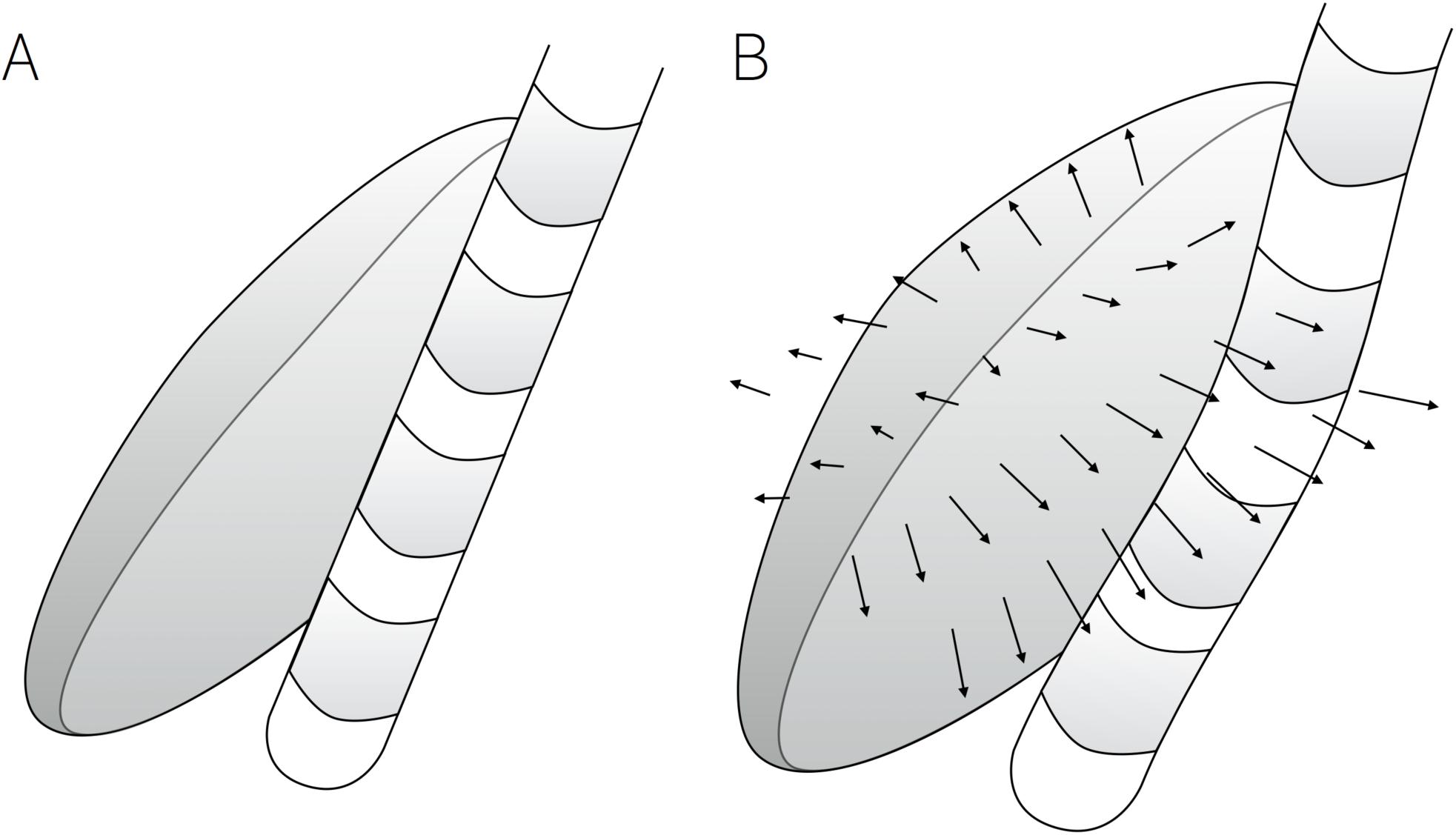
Relationship of a DBS electrode and the STN of a hypothetical patient in native single subject space (A) and standardized stereotactic (MNI) space (B). In standard space, the STN is defined once and for all based on the particular template. Arrows exemplarily illustrate the deformation field estimated to warp the (smaller) STN of the patient into the (larger) STN defined by the template. Note that this deformation leads to a bend in the DBS electrode and may also break up the distribution of electrode contacts (placed equidistantly on a straight line in A and on a curve with heterogeneous distances in B).

**Figure S2:**
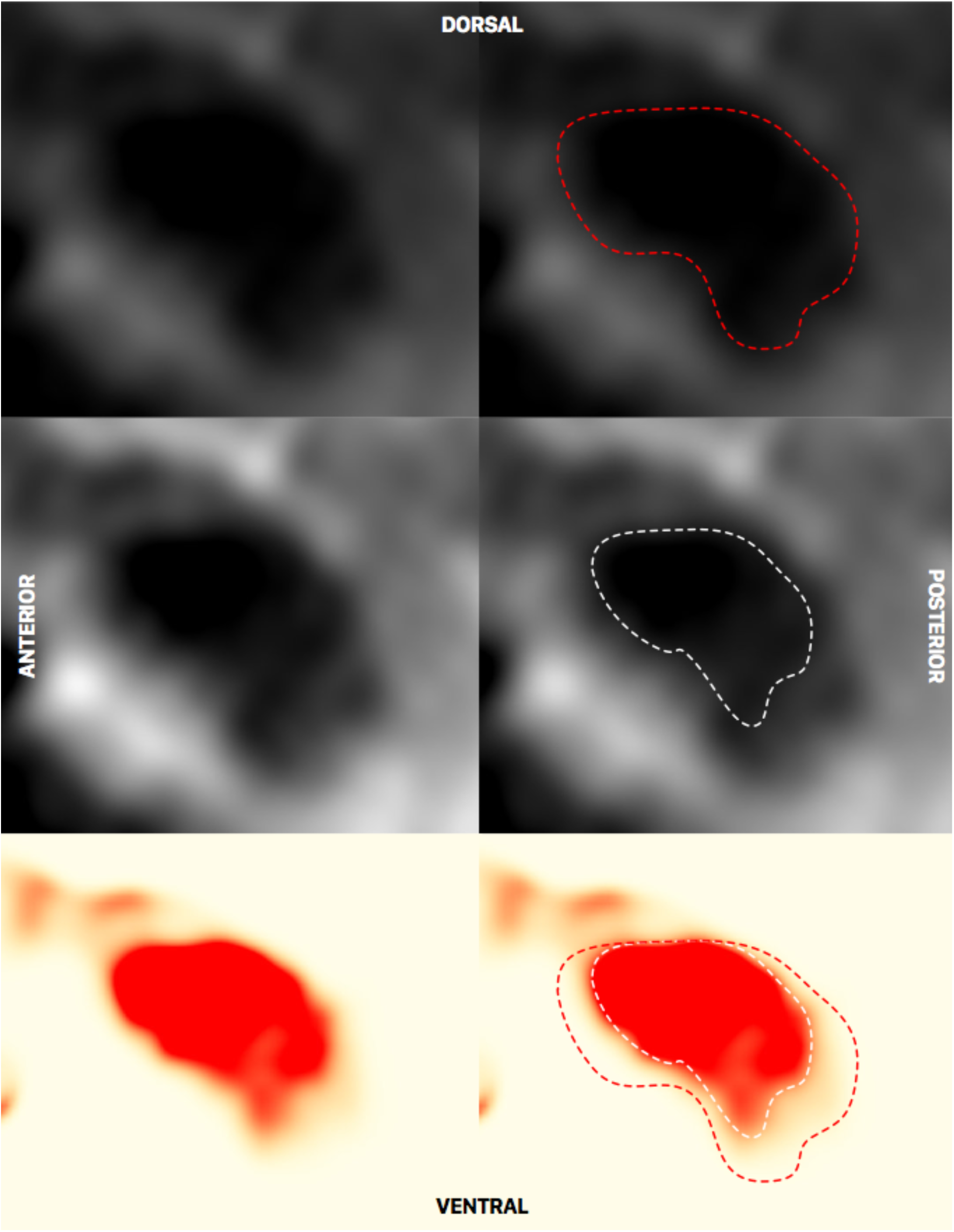
Sagittal slice of the T2-weighted ICBM template at x = -7.3 mm. Here, depending on windowing, the RN appears substantially larger (top) or smaller (middle). The RN second-level probability map (bottom) exhibits a much higher signal-to-noise ratio and is helpful in determining the ventral anatomical border of the nucleus in the process of manual segmentation. Based on neighboring slices, the correctness of the map could be confirmed to result from a partial volume effect.

